# Association of object mnemonic discrimination deficit with age and AD biomarkers in older adults

**DOI:** 10.64898/2026.09.14.750966

**Authors:** Zhengshi Yang, Xiaowei Zhuang, Lynn M. Bekris, Maria Khrestian, Katherine A. Koenig, Tim Curran, Rajesh Nandy, Pavithran Pattiam Giriprakash, Chendi Han, Jagan Pillai, Robert Fox, Mark J. Lowe, Dietmar Cordes

## Abstract

**Background:** Blood–based Alzheimer’s disease (AD) biomarkers provide scalable detection of AD pathology, yet their relationships with hippocampal–dependent mnemonic discrimination remain unclear. This study examined how object mnemonic discrimination relates to age, hippocampal structure, and plasma AD biomarkers in cognitively normal (CN) older adults and individuals with amnestic mild cognitive impairment (aMCI).

**Methods:** Forty–six CN and forty–one aMCI participants completed an object mnemonic similarity task (MST). Mnemonic discrimination was quantified using the overall lure discrimination index (LDI) and similarity–specific LDIs. Plasma pTau181, pTau217, pTau231, and GFAP were assayed, and hippocampal volumes were derived from 7T MR scanner. Multiple linear regression models assessed associations between LDI and biomarkers, controlling for age, sex, education, APOE4 status, and diagnosis. Repeated–measures ANCOVA and linear mixed effects models evaluated similarity–dependent age effects. ROC analyses and stepwise logistic regression quantified diagnostic classification performance.

**Results:** LDI was significantly reduced in aMCI compared to CN. Total hippocampal volume and plasma pTau217 showed the strongest associations with LDI, exceeding those of pTau181, pTau231, and GFAP. LDI discriminated aMCI from CN with high accuracy (AUC = 0.836), comparable to hippocampal volume and plasma biomarkers. A combined model including LDI, pTau181, pTau217, and hippocampal volume achieved the highest diagnostic accuracy (AUC = 0.931). Age effects were similarity–dependent: discrimination declined with age only for low–similarity lures, independent of diagnosis. Sex differences emerged within aMCI, with females showing lower LDI than males.

**Conclusions:** Object mnemonic discrimination provides a sensitive behavioral marker linked to hippocampal volume and plasma pTau217, showing discriminative power comparable to established AD biomarkers in separating aMCI from CN participants. Similarity–dependent age effects highlight age-related deficits are not universal for lures with varying similarity levels.

## BACKGROUND

Age-related memory decline is a common feature of normal aging and is further exacerbated in the early stages of Alzheimer’s disease (AD) [1]. Among the cognitive processes affected early in the disease, the ability to discriminate between similar experiences—often referred to as mnemonic discrimination or pattern separation—has received increasing attention [2, 3].

The mnemonic similarity task (MST) was specifically designed to assess object-based mnemonic discrimination and has been applied in both cognitively normal older adults and clinical populations [4]. Performance on the MST has been shown to decline with advancing age and to be disproportionately impaired in individuals with amnestic mild cognitive impairment (aMCI) [5]. Importantly, deficits on the MST may emerge even when traditional episodic memory measures remain relatively preserved [3]. Prior works suggest that hippocampal integrity is closely linked to mnemonic discrimination performance [4] In healthy aging, lower DG volume and higher CA3/DG activation are associated with worse lure discrimination, suggesting inefficient or compensatory hyperactivity [6]. Smaller medial temporal and hippocampal subfield volumes are associated with impaired mnemonic discrimination in MCI [7]. However, the extent to which mnemonic discrimination deficits relate to molecular biomarkers of AD is far from clear. Extensive evidence supports that blood-based biomarkers, including plasma phosphorylated tau (pTau) species and glial fibrillary acidic protein (GFAP), are reliable fluid biomarkers for AD assessment [8, 9]. The clinical relevance of object mnemonic discrimination performance in conjunction with hippocampal structural measures and plasma AD biomarkers remains to be tested. GFAP is a sensitive biomarker reflecting reactive astrocytosis and neuroinflammation that increases early in the AD continuum. Although GFAP is not directly linked to amyloid deposition or neurodegeneration, multiple studies have demonstrated that elevated plasma GFAP level is strongly associated with amyloid pathology and cognitive impairment, and clinical progression. Importantly, the diagnostic and prognostic performance of plasma GFAP has been shown to be comparable to that of plasma p–tau variants, supporting its relevance as a complementary marker of AD–related biological change.

In addition to age and biomarker burden, sex differences have been increasingly recognized in both cognitive aging and AD [10, 11]. Women show a higher lifetime risk of AD and may exhibit distinct cognitive trajectories during the prodromal stages of the disease [12, 13]. Nevertheless, sex-specific patterns in mnemonic discrimination performance in non-demented older adults have not been systematically examined.

In the present study, we investigated the association between object mnemonic discrimination deficits, age, hippocampal volume, and plasma AD biomarkers in a cohort of cognitively normal (CN) older adults and individuals with aMCI. We further examined whether mnemonic discrimination decline varies as a function of stimulus similarity and whether sex moderates the mnemonic discrimination deficits between aMCI and CN. By integrating behavioral, structural, and blood-based biomarker measures, this work aims to clarify the relative contributions of aging and AD-related pathology to hippocampal-dependent memory dysfunction in older adults.

## METHODS

### Subject Enrollment

One hundred participants (50 CN and 50 MCI) were initially recruited in the study from the Cleveland Clinic at Cleveland, Ohio since 2022. Before collecting MRI data, all participants underwent an outside-scanner training session and a mock run of our task. Only participants with a no-response rate less than 20%, and correct response rates greater than 60% for target and foil stimuli proceeded to MRI scan. The thresholds were heuristically chosen to ensure valid estimation of LDI. A total of thirteen participants (4 CN, 9 aMCI) were excluded prior to analysis. Exclusions were due to incomplete MRI acquisition or unusable MRI data (1 CN, 4 aMCI), inability to complete MRI because of claustrophobia (1 CN, 1 aMCI), and behavioral training failure (training accuracy below threshold or inability to complete the MST training; 2CN, 4 aMCI). These exclusion criteria were applied uniformly across diagnostic groups and reflect standard quality–control procedures for MRI acquisition and MST task performance. The exclusion rate due to behavioral training failure did not differ significantly between groups (χ²(1)=0.71, p=0.40; Fisher’s exact test p>0.05), suggesting that behavioral training failure was not disproportionately associated with diagnostic status and is unlikely to have introduced meaningful selection bias. A final cohort of 46 cognitively normal participants and 41 participants with aMCI were included in the analysis.

### Mnemonic Similarity Task

The objects used in this task and the similarity levels of lure objects were obtained from the original MST [14, 15]. Each participant underwent three consecutive 13-minute MST fMRI sessions with different sets of objects in a single visit. Each session consisted of an encoding phase, a memory recognition phase, and two distraction phases immediately after encoding and recognition phases. An instruction screen was presented for 5 seconds prior to each phase to remind subjects of the upcoming task. In the encoding phase, sixty-six everyday objects were shown sequentially for 3 seconds each on a white background. Participants indicated if each object was indoor or outdoor by pressing a button. In the recognition phase, eighty-six objects were shown sequentially at the center of the screen, including 22 targets, 44 lures and 20 foils. Lures differ in their resemblance to previously encoded objects, spanning high (17 objects), medium (16 objects), and low (11 objects) similarity levels. Participants decided if each object was the same, similar, or new compared to the encoding phase. In the distraction phase, participants judged whether two circles were equally dark, with 5 stimuli shown for 3 seconds each. More detailed description of this task can be found in our previous study [16]. Mnemonic discrimination performance was characterized by lure discrimination index (LDI) [4], which was calculated as the probability of identifying a lure as “similar” minus the probability of erroneously identifying a foil as “similar”. We also calculated the LDI for lures with high, medium, and low similarity levels separately, denoted as LDI_s_. LDI and the similarity-specific LDI_s_ were derived from lures aggregated across all three sessions, comprising 51 high-similarity, 48 medium-similarity, and 33 low-similarity trials. The reliability analysis suggested that similarity-specific measures exhibited good stability across sessions, supporting the robustness of the reported similarity-dependent effects (see Supplementary Material).

### Plasma AD biomarkers (pTau181, pTau217, pTau231, GFAP)

Plasma levels of pTau181 and pTau217 were measured using the S-PLEX Human Tau Kit from Meso Scale Discovery (K151APFS and K151AGMS, MSD; Rockville, MD), according to the manufacturer’s instructions. In short, an S-PLEX 96-Well SECTOR streptavidin plate was coated with a biotinylated capture antibody solution. Plasma samples were added to the plate followed by a TURBO-BOOST detection antibody solution, an S-PLEX enhance solution, and an S-PLEX detection solution. After all necessary incubations and washes, Read Buffer was added to the plate and it was immediately read on the MESO QuickPlex SQ 120MM instrument. The MSD Discovery Workbench software was used to analyze the raw data and calculate the concentration. The assay calibrators provided in the kit and the plasma samples were run neat in duplicate.

Plasma levels of GFAP and pTau231 were measured using commercially available digital immunoassays (Neuro 4-Plex E Advantage PLUS Kit and pTau231 Advantage PLUS Kit) on the Simoa SR-X platform (104465 and 104512, Quanterix; Billerica, MA) according to the manufacturer’s instructions. Briefly, magnetic beads coupled with a capture antibody are added to the sample, followed by target specific biotinylated detection antibodies and a streptavidin reporter enzyme conjugate. After this sample preparation is completed, the assay plate is inserted into the SR-X instrument. The instrument performs the run and produces analysis results within two hours. The assay calibrators provided in the kit were run neat in triplicate while control samples and plasma samples were diluted 1:4 for the N4PE kit and 1:2 for the pTau231 kit and run in duplicate.

### MRI Acquisition

The imaging data were acquired on a 7T Siemens MAGNETOM Terra MR scanner, using a 32-receive channel/8 transmit channel head coil at the Mellen Center of the Cleveland Clinic, Cleveland, Ohio. Standard B1 shimming without parallel RF transmission was used. Whole-brain T1-weighted 3D MP2-RAGE image, high-resolution hippocampus 3D T2-weighted image and MST fMRI data were acquired from each participant. Detailed acquisition protocol can be found in our previous study [16].

### Imaging biomarkers

FreeSurfer software suite (v7.2, https://surfer.nmr.mgh.harvard.edu/) was used to derive hippocampal volume. The T1 structural image was first analyzed with the main FreeSurfer command “recon-all” with default settings, and then the command “segmentHA_T2.sh” was used to segment hippocampal regions with T2 image fed into the analysis. The total hippocampal volume (left and right hemisphere combined) was extracted from the analysis. In addition, due to prior work linking dentate gyrus (DG) and CA3 dysfunction to pattern separation deficits [17], the total volume for CA3/DG was also computed. To account for individual differences in head size, the estimated total intracranial volume (eTIV) was used to normalize the absolute volumes. The relative volume, defined as 1000*(absolute volume/eTIV), was used in the following statistical analysis, where 1000 is an arbitrary factor.

### Statistical Analysis

Before conducting statistical analysis, we carried out a skewness test on plasma biomarkers and hippocampal volume to detect skewed data. All plasma pTau variants were highly skewed (skewness > 1) but not for plasma GFAP and hippocampal volume. A natural logarithm transformation was applied on the skewed measures before they entered the linear regression model. The skewness for plasma pTau181 and pTau217 after log transformation were 0.09 and 0.36, respectively. Plasma pTau231 was less skewed after log transformation, but it remained to be highly skewed (pre-transformation skewness = 8.00; post-transformation skewness = 1.11). The z-score of the transformed data was computed. The observations with |z|>3 were flagged as outliers, and they were winsorized by capping z-scores at ±3. GFAP and pTau231 each had a single outlier, and no outliers were identified for any other variables.

Multiple linear regression analysis was used to evaluate the association between LDI and plasma biomarkers or hippocampal volume, where age, sex, education, *APOE4* status, and clinical diagnosis were included as covariates. A base model including only covariates was first fitted, and the adjusted R² was calculated. We then evaluated the interaction between sex and diagnosis by adding a sex × diagnosis interaction term to the base model. Next, each plasma biomarker and hippocampal volume measure was added individually to the base model in separate analyses. Incremental model performance was quantified using ΔR², calculated as the difference in adjusted R² between the extended model and the base model. The standardized beta coefficient (β) and its 95% confidence interval derived from bootstrapping algorithm (N = 1000) were reported in the study. Bonferroni correction was used to correct for multiple comparisons. Influence and residual diagnostics suggested that the regression models were stable, well–behaved, and not unduly influenced by high–impact observations (see Supplementary Material).

To determine which factor had an interaction effect with lure similarity levels, a repeated-measures analysis of covariance (ANCOVA) model was applied to test if LDI_s_ at various lure similarity levels differs by diagnosis, age, sex, education, or *APOE4* status. Age was found to be the only significant factor in the analysis (see Results). Then we ran a linear mixed effect (LME) model to evaluate the differentiated age-effect. The LME model was formulated as LDI_s_ ∼ age*similarity + sex + education + *APOE4* + diagnosis + (1|subject), where age*similarity was the interaction effect, sex, education, *APOE4* status and diagnosis were covariates; and (1|subject) was the intra-subject random effect. We then computed the age slope at each lure level from fixed effects and evaluated its significance level based on Student’s *t* distribution.

Finally, we tested the prediction accuracy of LDI for diagnostic classification, in comparison with plasma AD biomarkers and hippocampal volume. The receiver operating characteristic (ROC) was used, and the area under the ROC curve (AUC) was computed for each predictor separately. To determine the best combination of these predictors to predict clinical diagnosis, a stepwise logistic regression algorithm was applied to select the optimal model with Akaike information criterion (AIC) as the optimization criterion. To assess multicollinearity, we computed variance inflation factors (VIFs) for all predictors retained in the final model. A DeLong’s test was carried out to evaluate if the optimal combined model was better than the best single predictor model. Harrell’s bootstrap optimism correction (1,000 iterations) was conducted to estimate the difference between apparent AUC in bootstrap samples and test–set AUC when applied to the original dataset. The optimism–corrected AUC was obtained by subtracting mean optimism from the apparent AUC. Calibration was evaluated using the Hosmer–Lemeshow goodness–of–fit test by comparing observed versus predicted probabilities across quintiles of predicted risk. Following STARD guidelines, we derived diagnostic accuracy metrics using the optimal probability threshold determined by Youden’s J statistic. We report sensitivity, specificity, positive predictive value (PPV), negative predictive value (NPV), and the confusion matrix for the internally validated model. All statistical analysis was performed in MATLAB R2025b (MathWorks, Natick, MA).

### Sensitivity Analysis

The years of education was significantly less in aMCI group compared to CN group (see *Results*). Because education is a well–established modifier of cognitive performance and may buffer against age–or pathology–related decline (reference), a sensitivity analysis was conducted to test if the primary findings were driven by the education difference between diagnostic groups. A one-to-one education–matching procedure was implemented to create a subsample in which CN and aMCI participants were comparable in their years of education. Using a nearest–neighbor procedure, each aMCI participant was matched to a CN participant whose education level was the closest and differed by no more than two years, with matching repeated sequentially until no additional eligible pairs could be found. The full analytical model was then re–estimated on this matched dataset.

## RESULTS

### Participants

The demographic characteristics, behavior data, and plasma biomarker levels in CN and aMCI participants were summarized in Table 1. Kruskal-Wallis tests were used to compare continuous variables, and the chi-squared tests were used to compare categorical variables between groups. These two groups did not show difference in age and sex, but the aMCI group had a higher proportion of *APOE4* carriers (51.2% for CN and 72.5% for aMCI), lower education, and worse LDI. Three CN and one aMCI participant did not have *APOE4* status available, and they were excluded from the following statistical analysis. The participants with aMCI showed significantly elevated plasma pTau181, pTau217, pTau231 and GFAP levels compared to CN. As to the task performance, the correct response rates for aMCI group were 66.9%, 37.9%, and 53.7% for target, lure, and foil, respectively. The correct response rates for CN group were 82.8%, 58.8% and 77.0% for target, lure, and foil, respectively. The CN group consistently showed better task performance than aMCI group across all three types of stimuli.

**Table 1.** Sample characteristics.

|  | CN | aMCI | pValue |
| --- | --- | --- | --- |
| Sex, male/female | 18 / 28 | 19 / 22 | 0.497 |
| APOE4<sup>a</sup>, carrier/non-carrier | 22 / 21 | 29 / 11 | 0.046 |
| Race, white/non-white | 42 / 4 | 40 / 1 | 0.210 |
| Age | 71 (68–74) | 72.00 (66–76) | 0.686 |
| Education | 17.50 (16.00–19.00) | 16.00 (12.75–18.00) | 0.018 |
| LDI | 0.40 (0.25–0.59) | 0.07 (-0.01–0.24) | 6.84e-08 |
| pTau181(fg/ml) | 926.6 (689.5–1164.9) | 1559.6 (1128.4–2072.8) | 1.43e-06 |
| pTau217(fg/ml) | 3627.6 (2740.0–4626.9) | 11389.4 (8241.3–15949.7) | 1.52e-09 |
| pTau231(pg/ml) | 3.51 (2.76–5.08) | 4.76 (3.65–6.31) | 8.92e-03 |
| GFAP (pg/ml) | 96.9 (71.4–124.3) | 140.5 (111.6–191.7) | 1.86e-05 |

Continuous measures are presented as median (interquartile range). P-values were derived from Kruskal-Wallis tests for continuous measures and chi-squared tests for categorical measures. CN = cognitively normal; aMCI = amnesic mild cognitive impairment.
<sup>a</sup>APOE4 status were unavailable for three CN participants and one aMCI participant.

### Contribution of Covariates in Predicting LDI

In the base model, clinical diagnosis (t = 5.97, p = 6.9x10^-8^) emerged as the sole factor significantly associated with LDI. Age, sex, education, and *APOE4* status did not demonstrate significant relationships with LDI. The base model had an adjusted R^2^ value of 0.355. When the interaction term between sex and diagnosis was included in the model, it showed a trend toward significance (p = 0.07; β = 15.91 95% CI [-1.35, 33.19]). Considering robust evidence for sex effects in AD, we additionally examined sex–specific simple effects using estimated marginal means with Tukey–Kramer correction as an exploratory follow–up analysis. These post-hoc comparisons revealed that females exhibited significantly lower LDI performance than males within the aMCI group (p = 0.035), consistent with the medium-to-high effect size (Cohen’s d = –0.65) identified with the raw score. No sex difference was observed in the CN group (p = 0.68; Cohen’s d = 0.15).

### Differentiated associations between LDI and AD biomarkers

When AD biomarkers were added to the base model, the β coefficients for the total hippocampal volume, pTau217, and pTau181, were observed to be significant after Bonferroni correction over the number of AD biomarkers considered (see Table 2). The model with total hippocampal volume had the strongest statistical significance with ΔR^2^ = 0.171, β = 10.56±2.38, p = 3.37x10^-5^. Plasma pTau217 had the strongest association among the fluid AD biomarkers with ΔR^2^= 0.112, β = -11.06±2.78, p = 1.62x10^-4^. The values in the plot were the residual after adjusting for age, sex, *APOE4* status, and education. The ΔR^2^ for plasma pTau181 was slightly lower than plasma pTau217 with ΔR^2^ = 0.089, β = -9.43±2.49, p = 3.11x10^-4^. The scatter plots, together with the fitting curves, between LDI and hippocampal volume, plasma pTau181 and plasma pTau217 were shown in Figure 2. The β coefficient for plasma pTau231 (uncorrected p = 0.024) and CA3DG volume (uncorrected p = 0.023) did not pass the Bonferroni correction, and the coefficient for plasma GFAP was not significant (p > 0.05). When we included the interaction term of these biomarkers with clinical diagnosis in the model, none of them were significant. The strongest interaction term was observed with plasma pTau231 (uncorrected p = 0.08). Due to the lower association of CA3DG volume compared to total hippocampal volume, CA3DG volume was not included in the further analysis.

**Figure 1.**
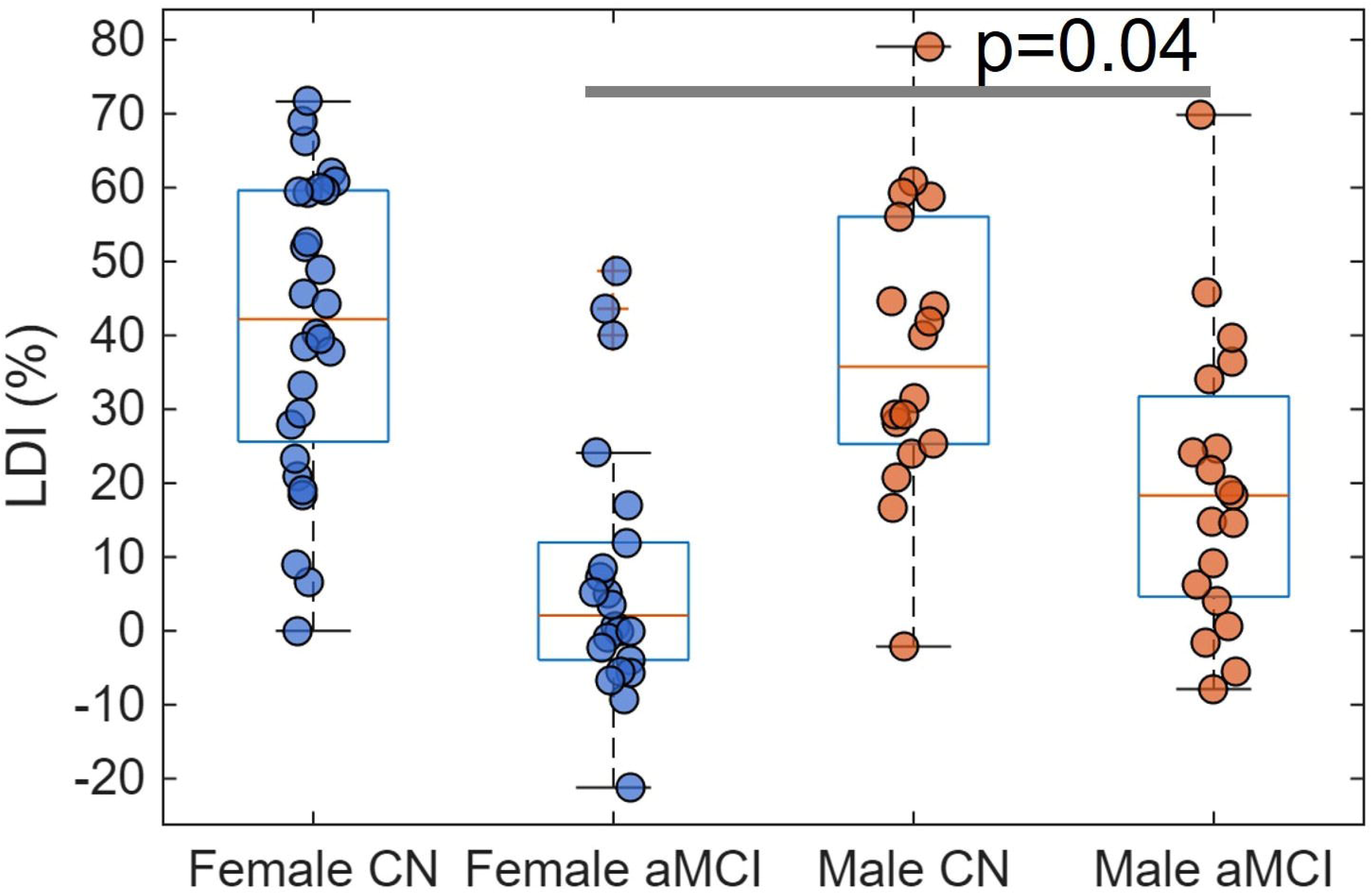
Boxplot of LDI grouped by sex and clinical diagnosis.

**Figure 2.**
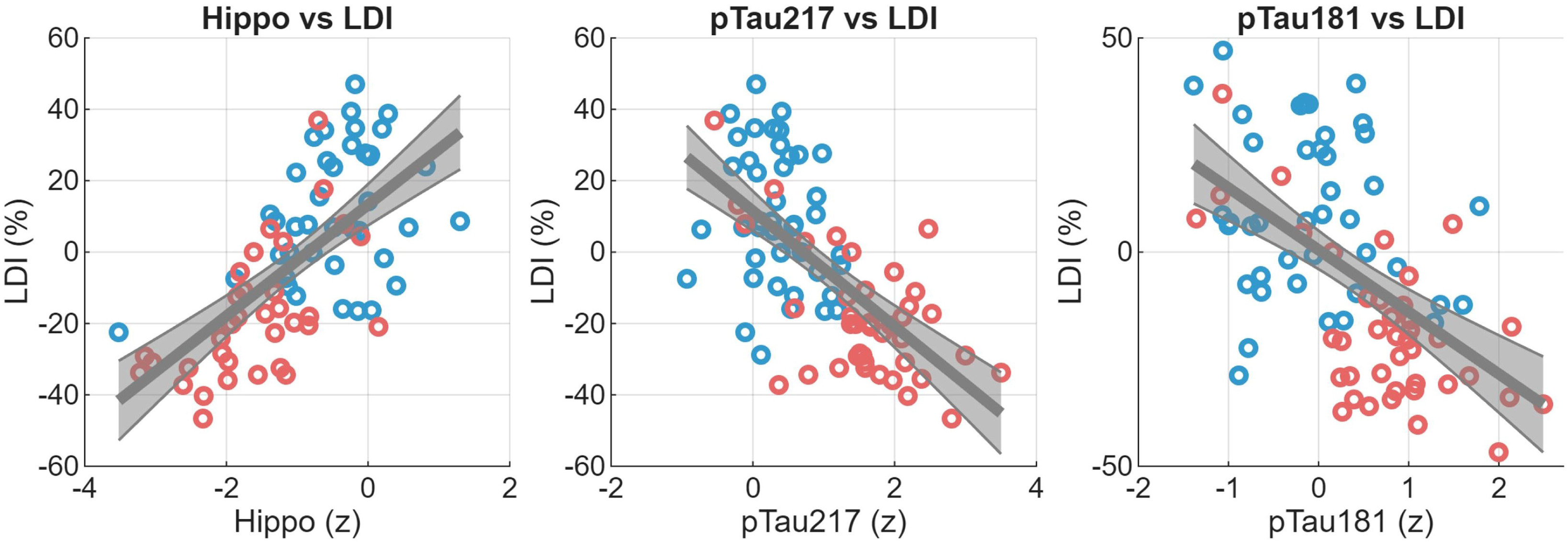
Scatter plots between LDI and AD biomarkers. The linear fitting analysis was carried out with subjects from both CN (blue circle) and aMCI (red circle) groups. Only the AD biomarkers showing significant association were shown in the figure. AD biomarkers were standardized. The residual of LDI and AD biomarkers after controlling age, sex, *APOE4* status, and education were used in the figure.

**Table 2.** The predictive power of AD biomarkers in predicting LDI.

| Row | Estimate | SE | t value | p value | 95% CI | $\Delta R^2$ |
| --- | --- | --- | --- | --- | --- | --- |
| (Intercept) | 42.37 | 35.15 | 1.21 | 0.23 | Not applied | Base model<br>adjusted $R^2$<br>= 0.355 |
| Age | -0.31 | 0.43 | -0.70 | 0.48 |  |  |
| Sex | 5.09 | 4.35 | 1.17 | 0.24 |  |  |
| Education | 0.97 | 0.80 | 1.21 | 0.23 |  |  |
| *APOE4* | 1.65 | 4.57 | 0.36 | 0.72 |  |  |
| Diagnosis | -27.32 | 4.58 | -5.97 | 6.92E-08 |  |  |
| pTau181 (z) | -9.43 | 2.49 | -3.78 | 0.00031 | [-14.67, -3.60] | 0.089 |
| pTau217 (z) | -11.06 | 2.78 | -3.97 | 0.00016 | [-17.27, -4.63] | 0.112 |
| pTau231 (z) | -2.84 | 2.17 | -1.31 | 0.195 | [-8.47, 2.42] | 0.006 |
| GFAP (z) | -5.82 | 2.52 | -2.31 | 0.024 | [-11.69, -0.35] | 0.034 |
| Hippo (z) | 10.56 | 2.38 | 4.44 | 3.37E-05 | [6.38, 14.87] | 0.171 |
| CA3DG (z) | 6.12 | 2.63 | 2.33 | 0.023 | [0.64, 10.31] | 0.023 |

A base model was carried out with only covariates included in the analysis, including age, sex, education and *APOE4*. The adjusted $R^2$ value was computed for the base model. $\Delta R^2$ = The difference of adjusted $R^2$ with and without imaging/plasma biomarker included in the model.

### LDI in Diagnosis Classification

In the single-predictor classification analysis, plasma pTau217 achieved the highest AUC value (AUC = 0.885, 95%CI [0.800, 0.955]). LDI demonstrated the second highest discriminative power in separating aMCI from CN with AUC = 0.836 (95%CI [0.748, 0.921]), which was higher than the AUC values obtained with the hippocampal volume (AUC = 0.809, 95%CI [0.706, 0.899]), plasma pTau181 (AUC = 0.808, 95%CI [0.699, 0.896]), plasma GFAP (AUC = 0.772, 95%CI [0.664, 0.872]), and plasma pTau231 (AUC = 0.666, 95%CI [0.545, 0.778]). The ROC curves for these measures were shown in Figure 3. To evaluate whether LDI differed in diagnostic accuracy from established biomarkers, we compared AUCs using DeLong’s test for these ROC curves. LDI showed comparable performance with pTau217 (p = 0.55, z = 0.60) and pTau181 (p = 0.07, z = 1.84), and it was significantly better than pTau231(p = 0.0008, z = 3.35), GFAP (p = 0.04, z = 2.06) and hippocampal volume (p = 0.004, z = 2.86).

**Figure 3.**
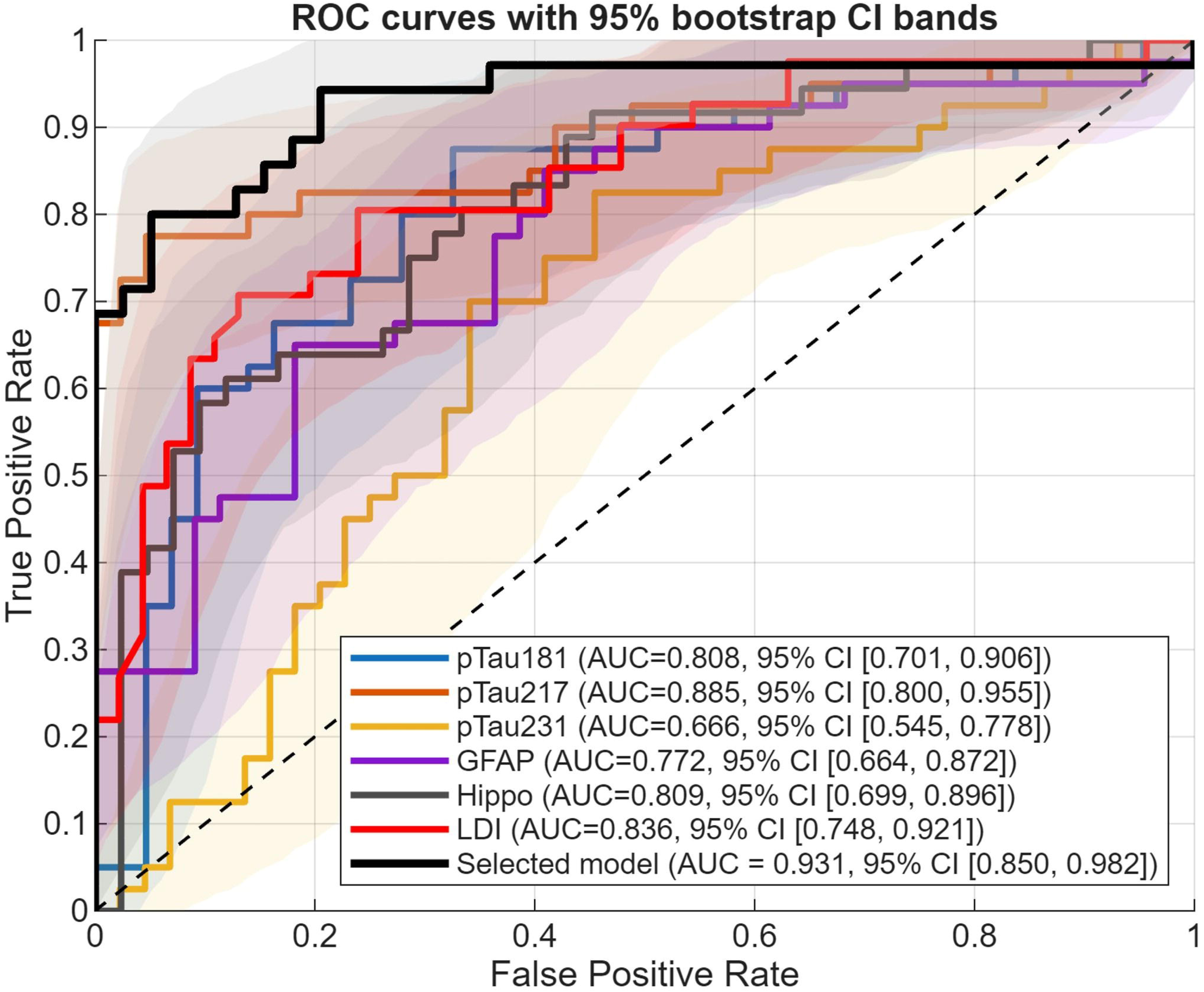
Receiver operating characteristics (ROC) curves for LDI, hippocampal volume, plasma biomarkers and the selected model in separating aMCI from CN. The shaded areas indicate the 95% confidence intervals for ROC curves derived from a bootstrapping algorithm.

In the stepwise logistic regression analysis, stepwise AIC selection identified a combined model including LDI (β = -1.157, 95%CI [-2.247, -0.067])), pTau181 (β = -0.995, 95%CI [-2.465, 0.474]), pTau217 (β = 2.453, [0.786, 4.121]), and hippocampal volume (β = -0.313, [-1.187, 0.561]). The variance inflation factors (VIF) for these predictors were low to moderate (LDI = 2.0, pTau181 = 3.6, pTau217 = 4.2, and hippocampal volume = 1.7). The AIC-selected model had an apparent AUC of 0.931, which was significantly better than the best single-predictor (pTau217) model with z = -2.352, p = 0.019. Bootstrap optimism correction yielded an optimism–adjusted AUC of 0.915 with 95% CI as [0.858, 0.991]. Calibration analysis demonstrated good agreement between predicted and observed probabilities. The Hosmer–Lemeshow test was nonsignificant (χ^2^=1.30, p = 0.73), suggesting that the model’s predictions well matched the actual observed data. Using the optimal threshold derived from Youden’s J (threshold = 0.66), sensitivity was 0.68, specificity 0.96, PPV 0.933, and NPV 0.772. The true positive / false positive / false negative / true positive at this threshold was 28/2/13/44.

### Similarity-dependent Age Effect on Mnemonic Discrimination Performance

Repeated-measures ANCOVA was carried out with the LDI_s_ measure to evaluate if age, sex, education, *APOE4* status, and diagnosis had differentiated influence on mnemonic discrimination performance depending on the lure similarity level. The LDI_s_ values significantly differed by clinical diagnosis (*p* = 7.0x10^-8^). The difference between clinical diagnosis was not differed by lure similarity level with the observation that the interaction term between diagnosis and similarity level was not significant (p = 0.14). The interaction between the intercept and lure similarity level was significant (p = 0.01), suggesting that the magnitude of LDI_s_ was differentiated by the lure similarity level. Age alone did not show association with LDI_s_ (p = 0.405), instead, the interaction term between age and lure similarity level was significant (p = 0.005), suggesting that age effect on LDI_s_ varied with the similarity level. All the other factors, including sex, *APOE4* status, and education, did not show significant main effect or interaction effect in the model. The 3-way interaction term between age, clinical diagnosis, and similarity level was not significant when it was included in the model, suggesting that the similarity-dependent age effect was not differentiated by clinical diagnosis.

In the linear mixed effect model, the high similarity group was treated as the reference group. The LDI_s_ for low similarity group was significantly differed from the reference group (p = 0.0002; Supplementary Table 1), such a difference was not observed for the medium similarity group (p > 0.05). There was no significant age effect on high similarity group (p > 0.05), and the age effect on the medium similarity group did not differ from the high similarity group. However, the age effect on the low similarity group was significantly different from the high similarity group (p = 0.0008). The linear fitting curves between age and similarity-dependent mnemonic discrimination performance, characterized by LDI_s_, are shown in Figure 4. The age slopes for high, medium, and low similarity lures were -0.073, -0.200, and -0.808, respectively. Only the slope for low similarity lures was significant (*p* = 0.04), and the slope for high (*p* = 0.87) and medium (*p* = 0.68) similarity lures were not significant.

**Figure 4.**
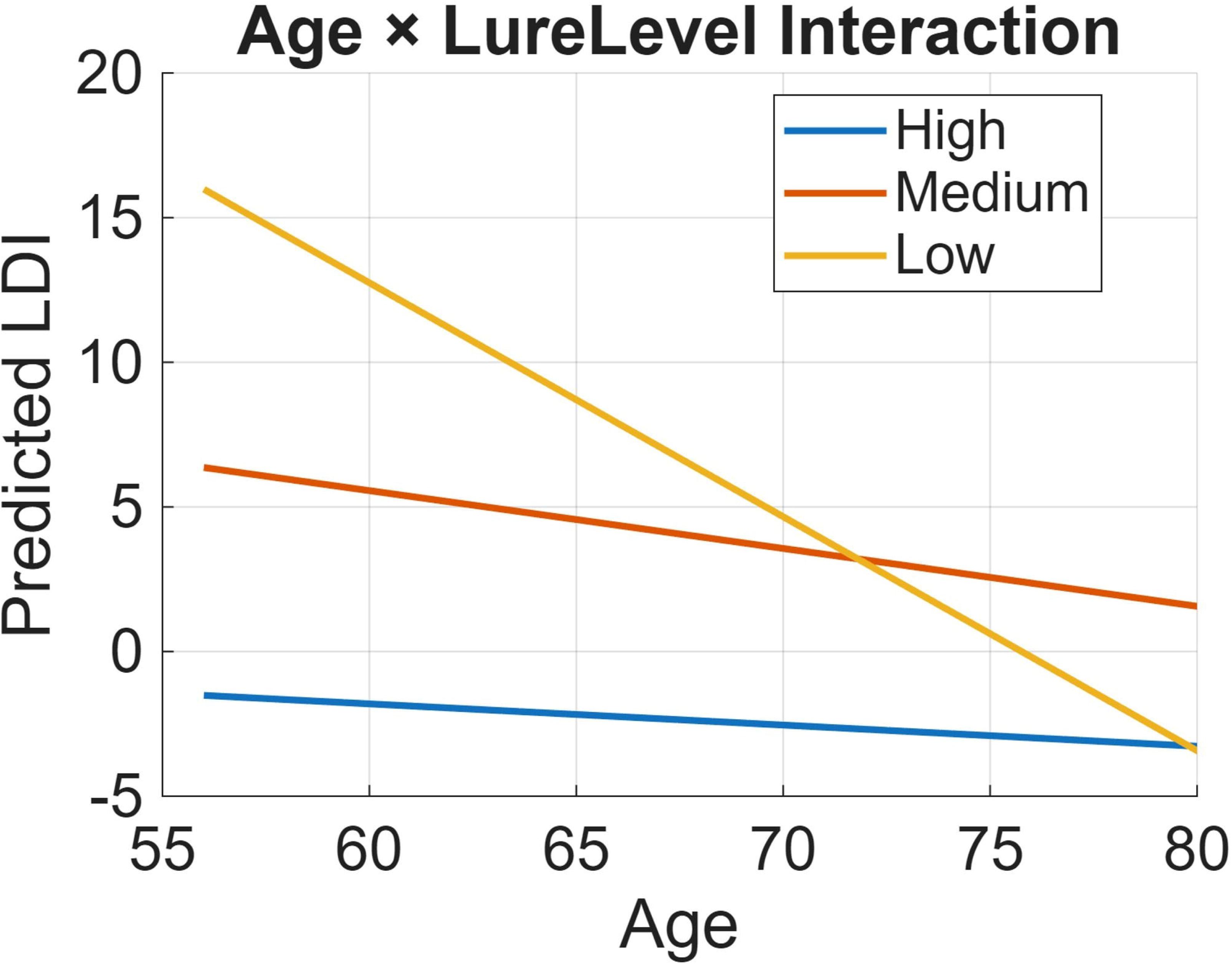
The similarity-dependent age effect on mnemonic discrimination performance. The fitting curves between LDIs and age for high, medium and low similarity lures were separately shown in the figure.

A total of 72 participants (36 CN and 36 aMCI) were included in the sensitivity analysis following the education-matching procedure. The years of education did not show difference between diagnostic groups in the matched sample (*p* = 0.57). We re-ran the full analytical analysis on this matched sample, The results with the subsample remained consistent with the findings observed with the full cohort. In fact, stronger sex-by-diagnosis interaction effect on LDI was observed with the subsample (p = 0.03) compared to the full cohort (p = 0.07).

## DISCUSSION

In this study, we investigated the relationships among object mnemonic discrimination, age, hippocampal volume, and plasma AD biomarkers among older adults. Three main findings emerged. First, overall mnemonic discrimination deficits were strongly associated with hippocampal volume followed by pTau217. Second, the discrimination capability for low–similarity lures declined with advancing age, whereas performance for high– and medium–similarity lures did not show a clear trend. Third, sex moderated mnemonic discrimination performance in aMCI but not in CN group.

### Hippocampal volume and plasma pTau217 as key correlates of mnemonic discrimination

Among all biomarkers examined, hippocampal volume showed the strongest association with LDI, accounting for the largest increase in explained variance beyond demographic and clinical covariates. This finding underscores the central role of hippocampal integrity in object mnemonic discrimination and is consistent with prior work linking hippocampal dysfunction to pattern separation deficits [17]. Although CA3/DG was of particular interest in pattern separation compared to other hippocampal subfields, the association was observed to be more robust for total hippocampal volume than the volume of CA3/DG. This finding seems to be counterintuitive, which could be because of multiple reasons. First, delineating hippocampal subfields is substantially more difficult than delineating the whole hippocampus due to the extremely small sizes of subfields, lack of contrast between them, and their complex structures [21]. These factors could compromise the reliability of subfield measures and weaken the statistical power in the analysis. Second, lesions in subfields other than CA3/DG could lead to mnemonic discrimination deficits [22]. CA1, together with CA3/DG, could act as a mismatch detector between previously encoded stimuli and the present lure stimuli [23]. This evidence suggests that CA3/DG is a critical area, but not the only area, in the hippocampus influencing mnemonic discrimination performance. Third, CA3/DG contributions to mnemonic discrimination might be more evident in functional measures than anatomical measures. Significant association between LDI and brain activation in left anterior CA3/DG was observed in our prior work [16].

Plasma pTau217 was more strongly associated with LDI than plasma pTau181, pTau231, and GFAP. This observation aligns with growing evidence that plasma pTau217 more closely reflects AD pathology, including amyloid, tau, and neurodegeneration, than other plasma pTau species [24]. Prior work demonstrated that plasma pTau217 was correlated to cognition and brain atrophy even in a cognitively healthy community cohort [25], which could explain why the association between pTau217 and LDI was not differentiated by clinical diagnosis. Briefly, the robust association between pTau217 and LDI suggests the potential utility of MST contributing to non-invasive cognitive assessment tool for AD screening.

### Contribution of education in MST

Education was not significantly associated with LDI performance in this cohort of older adults. Education is an important contributor to cognitive reserve, and higher educational attainment is associated with better performance and slower decline in multiple cognitive domains such as language, executive function, and episodic recall. However, to our knowledge, no published studies directly examined whether educational attainment was associated with mnemonic discrimination performance. Instead, a previous study showed that the performance in mnemonic similarity task was not significantly associated with long-term memory or executive function. These evidence collectively suggested that the influence of education might be weaker for tasks that require fine-grained perceptual discrimination heavily relied on hippocampus. Additionally, the relatively restricted range of education in our sample may have limited our ability to detect the association. Although the aMCI group was less educated than the CN group in our cohort, the sensitivity analysis demonstrated that the findings observed with the full cohort was not driven by the education difference between diagnostic groups.

### Sex differences in mnemonic discrimination in aMCI

In the exploratory analysis, female showed significantly worse mnemonic discrimination performance than males in aMCI group but not in CN group. This result is in line with steeper cognitive decline or greater vulnerability in female during the prodromal stages of AD.[26] Potential mechanisms include more rapid tau accumulation [27] and greater tau burden in female [28], hormonal changes associated with aging [29], or differential cognitive reserve [30]. Although the present study was not designed to directly test these mechanisms, the observed interaction highlights the importance of considering sex as a moderating factor in studies of cognitive aging and AD biomarkers.

### Diagnostic classification and clinical relevance

In the diagnostic classification, LDI demonstrated the second–highest discriminative performance, surpassed only by plasma pTau217, and exceeded the classification accuracy of hippocampal volume and several other plasma biomarkers. Behavioral assessments such as the MST are noninvasive, cost–effective, and easily deployable, making them attractive tools for large–scale screening and longitudinal monitoring. The stepwise logistic regression algorithm showed that a combination of LDI with pTau181, pTau217 and hippocampal volume was the best model for classifying diagnostic groups, suggesting that LDI may provide complementary information to these blood–based and imaging biomarkers. The VIFs for the four predictors in the best model fall within the commonly accepted range indicating low to moderate collinearity (VIF < 5), and none approached levels associated with model instability (VIF > 10). The opposite–signed β–weights for pTau181 and pTau217 therefore reflect their differential contributions after adjusting for shared variance rather than problematic multicollinearity. Taken together, the VIF results indicate that the stepwise model is stable and that inclusion of both pTau variants does not compromise interpretability.

### Age– and similarity–dependent effects on mnemonic discrimination

We did not find a significant age effect on the overall mnemonic discrimination performance, in contrast to worse LDI value in older adults than in younger adults in previous studies [18–20]. The discrepancy could be because participants were limited to older adults in our cohort. Instead, age effect was observed to depend on lure similarity levels. Specifically, discrimination of low–similarity lures declined significantly with age, whereas discrimination of highly similar lures did not show a measurable age–related decline. This pattern suggests that aging does not uniformly degrade mnemonic discrimination, but rather selectively affects processes engaged when discrimination demands are relatively low.

One possible reason is that discrimination of highly similar lures may already be low in older adults, limiting the sensitivity to detect further age–related decline. Alternatively, age–related changes in attentional allocation or decision criteria may disproportionately affect conditions in which perceptual overlap is less salient. Importantly, the similarity–dependent age effect was independent of clinical diagnosis, indicating that aging and AD pathology exert partially dissociable influences on mnemonic discrimination.

### Limitations and future directions

Several limitations should be acknowledged in the study. First, the cross–sectional design precludes causal inferences regarding the temporal relationships among mnemonic discrimination decline, biomarker changes, and hippocampal atrophy. Longitudinal studies will be necessary to determine whether LDI predicts future cognitive decline or biomarker progression. Second, although plasma biomarkers offer important advantages, the absence of amyloid or tau PET data limits direct comparisons with established in vivo measures of AD pathology. Considering that aMCI is a heterogeneous clinical syndrome encompassing individuals with and without underlying Alzheimer’s pathology, future work should focus specifically on aMCI participants with confirmed amyloid and tau positivity. Establishing biomarker–defined aMCI cohorts is a critical step toward determining whether MST performance reliably indexes early AD–related memory dysfunction and toward validating MST as a potential tool for early AD detection. Third, the observed sex-by-diagnosis interaction should be interpreted cautiously, as this effect was exploratory rather than a prespecified primary hypothesis. Although the pattern is suggestive, confirmation in an independent cohort with a larger sample size is necessary before drawing firm conclusions about sex–related modulation of mnemonic discrimination in aMCI.

## CONCLUSIONS

In this cross-sectional study, object mnemonic discrimination deficits were associated with smaller hippocampal volume and higher plasma pTau217 concentration. Task performance also showed meaningful diagnostic utility, with LDI distinguishing clinically defined aMCI from CN at a level approaching that of plasma pTau217. The similarity–dependent age effect and the presence of sex differences within aMCI highlight the multifactorial nature of mnemonic discrimination decline in older adulthood. Overall, these findings indicate that MST–derived LDI provides complementary information to plasma and imaging biomarkers in characterizing early AD–related cognitive impairment, although longitudinal and externally validated studies are needed to confirm these results.

## Declarations

### Ethics approval and consent to participate

This study was approved by Cleveland Clinic Institutional Review Board. All participants have given written, informed consent for their participation. The subjects’ consent was obtained according to the Declaration of Helsinki.

### Funding

This research project was supported by the National Institute on Aging (R01AG071566, R01AG074392, P30AG072959) and the National Institute of General Medical Sciences (P20GM109025), Cleveland Clinic Keep Memory Alive Young Investigator Award, a private grant from Stacie and Chuck Matthewson, a private grant from Peter and Angela Dal Pezzo, and a private grant from Lynn and William Weidner.

### Competing interests

All authors do not have any conflict of interest to disclose.

### Availability of data and materials

The data used in the study can be requested by sending a research proposal to the principal investigator (PI) Dr. Dietmar Cordes by using your institutional email address. The research proposal should not exceed 1-page. Please provide a clear and informative title for your proposed research. Please briefly describe the overall rationale for your study and summarize the specific aims/hypotheses that you will test with the specific data elements you are requesting.

### Authors’ contributions

Conceptualization, Z.Y., X.Z., M.J.L., P.P., C.H., J.P., and D.C.; methodology, Z.Y., X.Z., R.N., and D.C.; formal analysis, Z.Y.; writing—original draft, Z.Y.; writing—review and editing, Z.Y., X.Z., M.K., K.A.K., T.C., P.P., C.H., M.J.L., J.P., R.F., R.N., and D.C.; visualization, Z.Y.; supervision, D.C.; Data collection, M.K., K.A.K., X.Z., and M.J.L..

## Acknowledgements

Not applicable.

## Notes

### Competing Interest Statement

The authors have declared no competing interest.

