## Supplementary Material for "Association of object mnemonic discrimination deficit with age and AD biomarkers in older adults"

**Supplementary Material: Influence and Residual Diagnostics**

To assess the stability of the single‑biomarker linear regression models predicting LDI, we conducted a comprehensive set of influence diagnostics for each biomarker. Diagnostics included Cook’s distance, DFBETAS, and Studentized residuals. For each model, we summarized the maximum values of these diagnostics as well as the number of observations exceeding conservative screening thresholds.

Across all biomarker models, maximum Cook’s distance values ranged from 0.107 to 0.135, well below conventional thresholds for influential cases (Cook’s D > 1.0). Maximum DFBETAS values across all predictors ranged from 0.546 to 0.751, and biomarker‑specific DFBETAS ranged from 0.387 to 0.751, with the highest values observed for pTau217 and GFAP. Importantly, no observations exceeded the conventional influence threshold (|DFBETAS| > 1.0) for any coefficient in any model, indicating that no single participant meaningfully altered the biomarker–LDI associations. Maximum Studentized residuals ranged from 2.56 to 2.98, values consistent with well‑behaved residual distributions and not indicative of problematic outliers.

Together, these diagnostics demonstrate that the regression models were stable, well‑behaved, and not unduly influenced by high‑impact observations. No subjects were excluded based on influence diagnostics, and robust regression methods were evaluated but deemed unnecessary given the absence of influential observations under the conventional threshold.

**Reliability analysis.** To assess the reliability of similarity-specific LDI measures, LDIs were calculated separately for each of the three task sessions as the proportion of "similar" responses to lures minus the proportion of "similar" responses to foils. Inter-session reliability was assessed using the Spearman-Brown correction based on pairwise correlations among the three session-specific LDI estimates. Reliability estimates were 0.859 for the high-similarity condition, 0.898 for the medium-similarity condition, and 0.807 for the low-similarity condition.
