## Supplementary Table 1 for "Association of object mnemonic discrimination deficit with age and AD biomarkers in older adults"

Supplementary Table 1. Linear mixed effect model LDI_s_ ~ Age * LureLevel + Sex + Education + APOE4 + Diagnosis

+ (1|subjectID)

| Name | Estimate | SE | tStat | DF | pValue | Lower | Upper |
| --- | --- | --- | --- | --- | --- | --- | --- |
| (Intercept) | 21.85 | 34.77 | 0.63 | 239 | 0.530 | -46.64 | 90.35 |
| Age | -0.07 | 0.43 | -0.17 | 239 | 0.866 | -0.93 | 0.78 |
| Sex_Male | 5.06 | 4.15 | 1.22 | 239 | 0.224 | -3.12 | 13.25 |
| Education | 0.98 | 0.76 | 1.28 | 239 | 0.202 | -0.53 | 2.48 |
| APOE4_Carrier | 1.70 | 4.37 | 0.39 | 239 | 0.698 | -6.92 | 10.31 |
| Diagnosis_MCI | -27.12 | 4.38 | -6.19 | 239 | 2.54E-09 | -35.74 | -18.49 |
| Lure_Medium | 14.97 | 15.43 | 0.97 | 239 | 0.3331 | -15.43 | 45.36 |
| Lure_Low | 58.68 | 15.43 | 3.80 | 239 | 0.0002 | 28.28 | 89.08 |
| Age:Lure_Medium | -0.13 | 0.22 | -0.58 | 239 | 0.5613 | -0.56 | 0.30 |
| Age:Lure_Low | -0.74 | 0.22 | -3.38 | 239 | 0.0008 | -1.16 | -0.31 |
